# Oxidized phospholipid oxPAPC induces a Th1-like phenotype in regulatory T cells and inhibits their protective function in atherosclerosis

**DOI:** 10.1101/2023.02.08.527689

**Authors:** Brenna D. Appleton, Sydney A. Palmer, Harrison P. Smith, Lilly E. Stephens, Amy S. Major

## Abstract

**Background:** Regulatory T cells (T_regs_) are protective in atherosclerosis but reduced during disease progression due to cell death and loss of stability. However, the mechanisms of T_reg_ dysfunction remain unknown. Oxidized phospholipids (oxPLs) are abundant in atherosclerosis and can activate innate immune cells, but there is limited information regarding their impact on T cells. Given T_reg_ loss during atherosclerosis progression and oxPL levels in the plaque microenvironment, we sought to determine whether oxidized 1-palmitoyl-2-arachidonoyl-sn-glycero-3-phosphocholine (oxPAPC), an oxPL associated with atherosclerotic plaques, alters T_reg_ differentiation and function.

**Methods:** Naïve CD4^+^ T cells were cultured under T_reg,_ Th1, and Th17 polarizing conditions with or without oxPAPC and assessed by flow cytometry. Gene expression in oxPAPC-treated T_regs_ was analyzed by bulk RNA sequencing. Functional studies of oxPAPC-induced T_regs_ were performed by co-culturing T_regs_ with CTV-labeled CD8^+^ cells *in vitro. In vivo* suppression of atherosclerosis was evaluated by adoptively transferring control or oxPAPC-treated T_regs_ to hyperlipidemic *Ldlr*^*-/-*^ mice.

**Results:** Compared to controls, oxPAPC-treated T_regs_ were less viable but expressed higher levels of the Th1-associated markers T-bet, CXCR3, and IFN-γ. Th1 and Th17 skewing cultures were unaltered by oxPAPC. IFN-γ is linked to T_reg_ instability, thus T_reg_ polarization experiments were repeated using *Ifngr1*^*-/-*^ CD4^+^ T cells. IFNγR1 deficiency did not improve cell viability in oxPAPC-treated T_regs_, however, T-bet and IFN-γ expression was not increased suggesting a role for IFN-γ signaling. OxPAPC-treated T_regs_ were less suppressive *in vitro*, and adoptive transfer studies in hyperlipidemic *Ldlr*^*-/-*^ mice showed that oxPAPC-induced T_regs_ possessed altered tissue homing and were insufficient to inhibit atherosclerosis progression.

**Conclusions:** OxPAPC elicits T_reg_-specific changes that induce a Th1-like phenotype dependent on IFN-γ signaling. This is biologically relevant as oxPAPC-treated T_regs_ are unable to reduce atherosclerosis progression in *Ldlr*^*-/-*^ mice. This study supports a role for oxPLs in negatively impacting T_reg_ differentiation and atheroprotective function.

## Introduction

Atherosclerosis, the most common form of cardiovascular disease, is characterized by the accumulation of oxidized lipids in the artery wall which can drive immune cell recruitment to the intima. Although macrophages are important in atherogenesis and comprise a large portion of the immune cells in plaques, recent single-cell RNA sequencing of patient carotid plaques demonstrated that T cells make up at least 50% of infiltrating immune cells^1,2^. The importance of T cells to the disease progress in animal models has been previously demonstrated by our group^3^ and others (reviewed in ^4^), but less is known about how T cells are impacted by the oxidized lipid-rich environment of the aorta.

Multiple CD4^+^ T cell populations have been shown to influence the progression of atherosclerosis (reviewed in ^4^). Interferon-γ (IFN-γ)-producing Th1 cells are generally considered atherogenic due to their ability to stimulate inflammatory cell recruitment to the intima, activate dendritic cells, and promote foam cell formation^5–8^. However, while IFN-γ deficiency is protective in male *Apoe*^*-/*-^ mice, female mice are not protected^9^, suggesting the role of Th1 cells and IFN-γ in atherosclerosis may be more complex. Effects of Th17 cells in atherosclerosis are even less concrete with several groups reporting either atherogenic or atheroprotective effects depending on the mouse models used or the disease stage studied^10–13^.

Ample evidence suggests regulatory T cells (T_regs_) are protective in atherosclerosis both through direct interactions with coinhibitory receptors and production of anti-inflammatory cytokines^14,15^. Depletion of T_regs_ from atherosclerosis susceptible mice significantly increases plaque size and progression^14,16^. Additionally, T_regs_ are increased in multiple mouse models of regressing plaques and are required for regression to take place^17^.

Studies in both mice and humans have demonstrated reduced T_reg_ numbers as atherosclerosis progresses^18–20^. This loss may be in part due to an increase in T_reg_ apoptosis^20,21^. T_regs_ in atherosclerosis lose stability and convert to exT_regs_, adopting the phenotype and function of other T helper subtypes like Th1 or T follicular helper cells, both of which are atherogenic rather than protective^22,23^. Each of these models provides a hypothesis for the overall reduction of T_regs_ during atherosclerosis, but mechanisms underpinning T_reg_ death and loss of stability in the plaque microenvironment remain unknown. Collectively, these findings highlight that T_regs_ are critical mediators for controlling and reversing atherosclerosis.

Oxidized low-density lipoprotein (oxLDL), one of the main oxidized lipoproteins associated with atherosclerosis, is highly abundant in the lesion environment and has been shown to both inhibit T_reg_ differentiation and promote apoptosis *in vitro*^21,22,24,25^. OxLDL is a complex particle made up of the central protein, apolipoprotein B100, and a variety of oxidized lipids, including the phospholipid, oxidized 1-palmitoyl-2-arachidonoyl-sn-glycero-3-phosphocholine (oxPAPC)^26^. Interestingly, several studies have demonstrated that oxPAPC is the bioactive component of oxLDL, eliciting many of the same effects in innate immune cells as oxLDL despite its reduced molecular size and complexity^27–29^. OxPAPC is also increased in the membrane of apoptotic cells which are abundant in atherosclerotic plaques^30^. However, the effects of oxPAPC have mostly been studied in innate immune cells and its direct impact on T cells is unknown.

In this study we sought to determine if oxPAPC affects T cell differentiation and function to promote atherosclerosis. We report that oxPAPC altered T_reg_ polarization, inducing T_regs_ to develop a Th1-like phenotype, expressing T-bet and producing IFN-γ. These effects were specific to T_regs_, as other T helper subtypes were unaffected. OxPAPC-treated T_regs_ had reduced suppressive activity *in vitro* and, when adoptively transferred into hyperlipidemic *Ldlr*^*-/-*^ mice, oxPAPC-treated T_regs_ did not inhibit plaque progression compared to control T_regs_. Overall, our findings demonstrate that oxPAPC alters T_reg_ phenotype and makes them less protective against plaque progression. Therefore, exposure of differentiating T_regs_ to oxPAPC may be an important mechanism for their dysfunction during atherosclerosis.

## Materials and Methods

### Mice

C57BL/6J (B6; Stock # 000664), B6.129S7-Ldlr^tm1Her^/J (*Ldlr*^*-/-*^; Stock # 002207), B6.129S7-Ifngr1^tm1Agt^/J (*Ifngr1*^*-/-*^; Stock # 003288), B6.Cg-Foxp3^tm2Tch^/J (B6-FoxP3^GFP^; Stock # 006772), and B6.PL-Thy1^a^/CyJ (Thy1.1; Stock # 000406) mice were originally obtained from the Jackson Laboratory (Bar Harbor, ME). B6-FoxP3^GFP^ and Thy1.1 animals were crossed to generate B6-Thy1.1-FoxP3^GFP^ mice. Animals were maintained and housed at Vanderbilt University. All mice used in these studies were on the C57BL/6J background. Procedures were approved by the Vanderbilt University Institutional Animal Care and Use Committee. Both male and female mice were used for studies as specified.

### T Cell Polarization

Spleens from four- to six-week-old mice were processed to single cell suspensions and CD4^+^ T cells were enriched using mouse CD4 (L3T4) Microbeads kit (Miltenyi Biotec Cat # 130-117-043) according to manufacturer instructions. Experiments were performed using both male and female mice. For T_reg_ polarization, CD4^+^ T cells were cultured in *α*-CD3 (coating concentration was 2*μ*g/ml; Tonbo Cat # 40-0031) coated flat-bottom 96-well plates at a final concentration of 2×10^6^ cells/ml in T Cell Media (TCM; RPMI-1640, 10% fetal bovine serum [FBS], L-glutamine-penicillin-streptomycin [2mM, 100 units, 100*μ*g/ml, respectively], 2mM β-mercaptoethanol, and 1x non-essential amino acids) with 2μg/ml α-CD28 (Tonbo Cat # 70-0281), 50U/ml IL-2 (Tonbo Cat # 21-8021 and Peprotech Cat # 212-12), 10ng/ml TGF-β (Peprotech Cat # 100-21), 2μg/ml α-IFN-γ (Tonbo Cat # 40-7311), and 1μg/ml α-IL-4 (Tonbo Cat # 70-7041) for three days. If culture was continued to day 5, T_regs_ were split, replated, and fed with TCM containing final concentrations of 50U/ml IL-2, 10ng/ml TGF-β, 2μg/ml α-IFN-γ, and 500ng/ml α-IL-4. Media was replaced on day 4. T_regs_ were cultured with or without 5μg/ml oxPAPC (Invivogen Cat # tlrl-oxp1) for the first three days of culture unless otherwise stated. For Th1 polarization, CD4^+^ T cells were cultured in α-CD3 coated flat-bottom 96-well plates at a final concentration of 2×10^6^ cells/ml in TCM with 2μg/ml α-CD28, 10ng/ml IL-12 (Peprotech Cat # 210-12), and 1μg/ml α-IL-4 for three days. On day 3, cells were split, replated, and fed with TCM containing final concentrations of 20U/ml IL-2, 10ng/ml IL-12, and 1μg/ml α-IL-4. Feeding was repeated on day 4. For Th17 polarization, CD4^+^ T cells were cultured in α-CD3 coated flat-bottom 96-well plates at a final concentration of 1×10^6^ cells/ml in TCM with 2μg/ml α-CD28, 25ng/ml IL-6 (Peprotech Cat # 216-16), 5ng/ml TGF-β, 2μg/ml α-IFN-γ, and 500ng/ml α-IL-4 for three days. Th1 and Th17 cells were cultured with or without 5μg/ml oxPAPC for the duration of the culture.

### Flow Cytometry

Flow cytometry was performed on cells obtained *ex vivo* and following *in vitro* cell culture. Cells requiring a viability dye were stained with Viobility Fixable Dye (Miltenyi Biotec) or Ghost Dyes™ (Tonbo) according to the manufacturer protocols. For surface staining, cells were washed in HBSS containing, 1% BSA, 4.17mM sodium bicarbonate, and 3.08mM sodium azide (FACS buffer), followed by a 10-minute room temperature incubation in 1μg/ml Fc block (α-CD16/32; Tonbo Cat # 40-0161) diluted in FACS buffer. Cells were then stained for 30 minutes at 4°C protected from light with the following antibodies diluted in FACS buffer: α-B220-APCCy7 (Tonbo Cat # 25-0452), α-CD4-PECy7 (Tonbo Cat # 60-0042), α-CD4-PerCp-Cy5.5 (Tonbo Cat # 65-0042), #), α-CD4-SB600 (eBioscience Cat # 63-0041-82), α-CD8a-APCCy7 (Tonbo Cat # 25-0081), α-CD11b-APCCy7 (BD Biosciences Cat # 557657), α-CD25-FITC (Tonbo Cat # 35-0251), α-CD45.2-PerCp-Cy5.5 (Tonbo Cat # 65-0454), α-CXCR3-APC (BD Biosciences Cat # 562266), α-GITR-FITC (Tonbo Cat # 35-5874), α-ICOS-FITC (eBioscience Cat # 11-9949-82), α-Thy1.1-V450 (BD Biosciences Cat # 561406). Samples not requiring intracellular staining were then washed in FACS buffer and fixed in 2% paraformaldehyde (PFA). Samples with intracellular stains were washed then permeabilized and stained with the following intracellular antibodies according to the FoxP3/Transcription Factor Staining Buffer Set manufacturer protocol (eBioscience Cat # 00-5523-00): α-FoxP3-PECy7 (eBioscience Cat # 25-5773-82), α-IFN-γ-APC (Tonbo Cat # 20-7311), α-IL-4-FITC (eBioscience Cat # 11-7042-82), α-IL-17-PE (BD Biosciences Cat # 559502), α-T-bet-PE (Biolegend Cat # 644810), α-ROR-γt-PerCp-Cy5.5 (eBioscience Cat # 12-6988-82). All samples were washed and resuspended in 2% PFA before analysis. Cells that were stimulated with phorbol myristate acetate (PMA; Sigma Cat # P1585) and ionomycin (Sigma Cat # I9657) prior to staining were treated with either 20ng/ml PMA, 1μg/ml ionomycin, and 1.3μg/ml Golgi Stop (BD Biosciences Cat # 554724) or 500ng/ml PMA, 500ng/ml ionomycin, and 110μg/ml Golgi Plug (BD Biosciences Cat # 51-2301KZ) in TCM or complete RPMI (cRPMI; RPMI-1640, 10% fetal bovine serum [FBS], L-glutamine-penicillin-streptomycin [2mM, 100 units, 100μg/ml, respectively], and 2mM β-mercaptoethanol) for four-six hours.

Apoptosis staining was done using the Annexin V Apoptosis Detection kit (Tonbo Cat # 20-6410) and performed according to manufacturer instruction. Sample acquisition was performed on a MacsQuant Analyzer (Miltenyi Biotec) and data were analyzed using FlowJo Single Cell Analysis software.

### RNA Sequencing (RNAseq)

4.5×10^6^ - 7.3×10^6^ live control or oxPAPC-treated T_regs_ were harvested on day 3 of culture and RNA was isolated using RNeasy Plus Mini Kit (Qiagen Cat #: 74134) according to manufacturer instruction. RNA was sent to Vanderbilt Technologies for Advanced Genomics (VANTAGE) core. Libraries were prepared using 200-500ng of total RNA using NEBNext® Poly(A) selection kit and sequenced at Paired-End 150bp on the Illumina NovaSeq 6000 targeting an average of 50M reads per sample. Additional analysis using the resulting demultiplexed FASTQ files containing PF reads was performed by Vanderbilt Technologies for Advanced Genomics Analysis and Research Design (VANGARD). Sequencing reads were aligned against the mouse GENCODE GRCm38.p6 using STAR software v2.7.8a. Mapped reads were assigned to gene features and quantified using featureCounts v2.0.2. Normalization and differential expression were performed using DESeq2 v1.30.1. Significantly differentially expressed genes (fold change ≥2 and FDR ≤0.05) were used for subsequent Gene Set Enrichment Analysis v4.2.2.

### Diet Comparison Studies

Eight-week-old male and female *Ldlr*^*-/-*^ mice were maintained on normal chow or placed on Western diet (21% saturated fat and 0.15% cholesterol; Envigo Cat #: TD.88137) for 16 weeks. After which animals were sacrificed, and aorta and spleen were collected for flow cytometry analysis.

### Tissue Collection

Aortas were processed as previously described^31^. Briefly, aortas were perfused with PBS, cleaned of fat, minced, and digested with 450U/ml Collagenase Type I (Worthington Biochemicals Cat # LS004194), 125U/ml Collagenase Type XI (Sigma Cat # C-7657), 60U/ml Hyaluronidase Type I (Sigma Cat # H-3506), and 60U/ml DNase I (Millipore Sigma Cat # 69182) in PBS at 37°C for 30-45 minutes. Digested aortas were passed through 70μm cell strainers, and leukocytes washed before use in flow cytometry.

Livers were processed as previously described^32^. Briefly, livers were perfused with PBS, minced, and then digested with 1mg/ml Collagenase Type II (Gibco Cat # 17101-015) in HBSS with calcium and magnesium at 37°C for 30 minutes. Liver tissue was passed through 70μm cell strainers and allowed to settle for 45 minutes on ice. Supernatants were pelleted and resuspended in cold 40% Percoll with a 60% Percoll underlay. Gradients were centrifuged at 2000rpm at 10°C for 20 minutes with no break. Leukocytes were collected from the gradient interface.

### T_reg_ Suppression Assay

T_regs_ were skewed as described above. On day 5 of culture, T_regs_ were harvested and replated in cRPMI in flat-bottom 96-well plates at ratios of 3:1, 2:1, 1:1, 1:2, 1:4, and 1:8 with CD8^+^ T cells enriched using mouse CD8 (Ly-2) Microbeads kit (Miltenyi Biotec Cat # 130-049-401) and labeled with CellTrace Violet (CTV; Invitrogen Cat # C34557), both according to the manufacturer protocols. Also included in the co-culture, were 2×10^5^ irradiated (30Gy) feeder splenocytes per well and 1μg/ml α-CD3. After 72 hours, T_reg_ suppression was determined by measuring CTV dilution in CD8^+^ responder cells via flow cytometry. Percent inhibition was calculated as [100%-((proliferation of given T_reg_:T_res_ ratio/proliferation of 0:1 T_reg_:T_res_ group)x100)].

### Adoptive Transfers

Adoptive transfer protocol was adapted from Li *et al*.^33^. 15-week-old female *Ldlr*^*-/-*^ mice were placed on Western diet for 9-11 weeks after which mice were retro-orbitally injected with saline, or 1×10^6^ day 5 control or oxPAPC-treated T_regs_. The injections were repeated 4-6 weeks later, and animals were sacrificed 2 weeks after the final injection. Aortas, aorta draining lymph nodes (adLNs), livers, blood, and spleens were collected for flow analysis. Atherosclerosis severity was quantified in the proximal aorta using oil-red-O staining of 10μm cryosections as previously described^34^.

### Cholesterol and Triglyceride Assays

Mice were fasted for four hours, and blood was collected from the retroorbital sinus and centrifuged to obtain serum. Cholesterol (Millipore Sigma Cat # MAK436-1KT) and triglycerides (Millipore Sigma Cat # MAK266-1KT) were measured using commercially available kits according to the manufacturer instructions.

### ELISAs

Anti-oxLDL ELISAs were performed as previously described^32^. Briefly, Nunc Maxisorp 96-well plates (Invitrogen Cat # 44-2404-21) were coated in 1μg/ml oxLDL overnight at 4°C. After washing with 0.05% Tween in PBS and blocking with 3% BSA/PBS, serum samples were applied, and the plate was again incubated overnight. Biotinylated α-IgG (eBioscience Cat # 13-4013-85), α-IgG1 (Southern Biotech Cat # 1070-08), α-IgG2c (Southern Biotech Cat # 1080-08), or α-IgM (Southern Biotech Cat # 1020-08) were incubated for one hour at room temperature, followed by streptavidin-peroxidase (Southern Biotech Cat # 7200-05) for 30 minutes at room temperature. Plates were washed and OptEIA TMB substrate (BD Biosciences Cat # 555214) was applied for 8, 10, 15, and 5 minutes, respectively, before reaction was quenched with 2M HCl. Results were read at 450nm.

### Statistical Analyses

Normally distributed data was analyzed using Student’s *t*-tests (for two groups) or one-way ANOVAs with Bonferroni post-test (for three or more groups). For experiments with two independent variables, two-way ANOVAs with a Tukey’s multiple comparison test was used. All statistical tests were performed using GraphPad Prism.

## Results

### OxPAPC alters T_reg_ phenotype but does not impact Th1 or Th17 polarization

T_regs_ are known to be atheroprotective but are reduced by cell death or loss of stability in atherosclerosis, presumably due to the plaque microenvironment^14,20,22,23^. Oxidized phospholipids (oxPLs) are an abundant component of the lesion environment and can impact the functioning of immune cells^29,35^. To determine whether the oxPLs associated with atherosclerotic lesions, such as oxPAPC, contribute to T_reg_ dysregulation, we performed *in vitro* CD4^+^ T cell polarization experiments with or without oxPAPC. Skewing T_regs_ in the presence of oxPAPC significantly reduced their viability at day 3 in culture by increasing apoptosis (Figure 1A and Supplemental Figures 1A and 2). The proportion of live cells that express FoxP3 was not affected by oxPAPC (Figure 1A, middle panels). Interestingly, there was an increased proportion of T-bet^+^ FoxP3^+^ cells in oxPAPC-treated cultures (Figure 1A, bottom panels and Supplemental Figure 1A. Both cell death and dysregulated transcription factor expression were specific to T_reg_ skewing conditions as oxPAPC treatment had no effect on Th1 or Th17 viability or polarization (Figure 1B-C and Supplemental Figure 1B-C). Interestingly, oxPAPC was required during differentiation of naïve CD4^+^ T cells to T_regs_ as addition of the phospholipid after day 3, when FoxP3 levels are established, had no effect on cell viability or T-bet expression (Figure 1D). Collectively, these data support oxPL-mediated dysregulation of CD4^+^ T cell differentiation in a T_reg_-specific manner.

**Figure 1:**
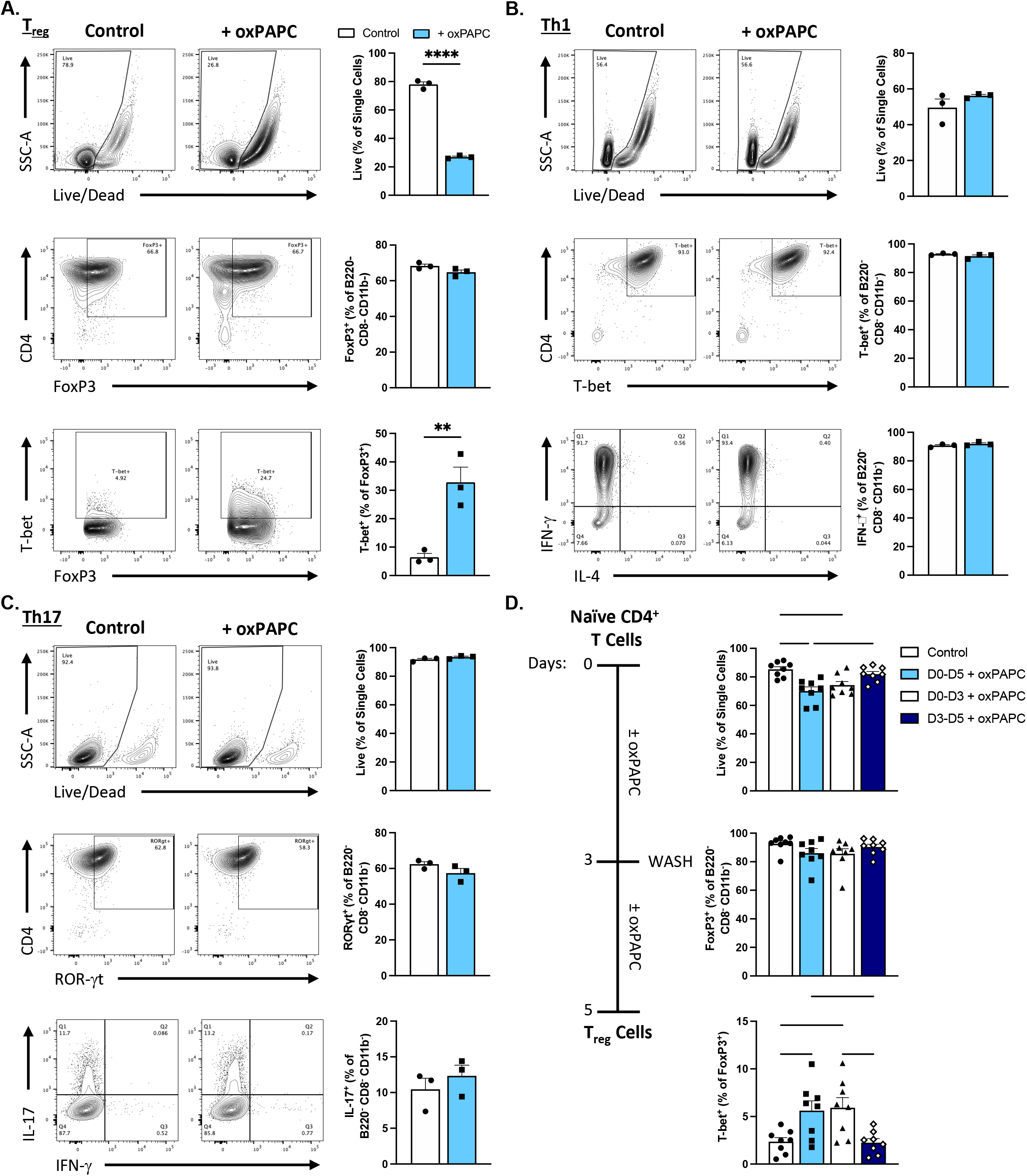
OxPAPC specifically alters T_reg_ phenotype. CD4^+^ T cells were enriched from the spleens of four- to six-week-old B6 mice and skewed to the indicated phenotype for three **(A and C)** or five **(B and D)** days with or without 5μg/ml oxPAPC. T cell phenotype was assessed by flow cytometry directly after harvest **(A**,**D)** or after a four-hour stimulation with PMA, ionomycin, and Golgi inhibitor **(B**,**C). (A-C)** Shown is one representative of at least three experiments.. ** and **** denote significance *p*<0.01 and *p*<0.0001, respectively, by Student’s *t* test. **(D)** CD4^+^ T cells were enriched from the spleens of four- to six-week-old B6 mice and skewed to a T_reg_ phenotype for five days with the indicated oxPAPC treatments. T_reg_ phenotype was assessed by flow cytometry directly after harvest *, **, and *** denote significance *p*<0.05, *p*<0.01, and *p*<0.001, respectively, by one-way ANOVA and Tukey’s multiple comparison. Data points represent individual mice and error bars show SEM.

In addition, though oxPAPC-treated T_regs_ had the same overall proportion of FoxP3^+^ cells as control T_regs_, the level of FoxP3 expression was altered. When oxPAPC-treated T_regs_ were analyzed on day 3, just following activation and differentiation, FoxP3 expression was significantly higher than in control T_regs_, and a greater proportion of the cells were FoxP3^hi^ (Supplemental Figure 3A-B). However, on day 5, after resting, oxPAPC-treated T_regs_ did not increase FoxP3 expression as controls did and a greater proportion of oxPAPC-induced T_regs_ were FoxP3^lo^ (Supplemental Figure 3A-B). T_reg_ functional markers (CD25, GITR, and ICOS) were concentrated in the FoxP3^hi^ group and Th1-like markers (T-bet, IFN-γ, CXCR3) were increased in the FoxP3^lo^ population (Supplemental Figure 3C). These data suggest that skewing T_regs_ in the presence of oxPAPC decreases FoxP3 stability resulting in increased T effector function.

**Figure 2:**
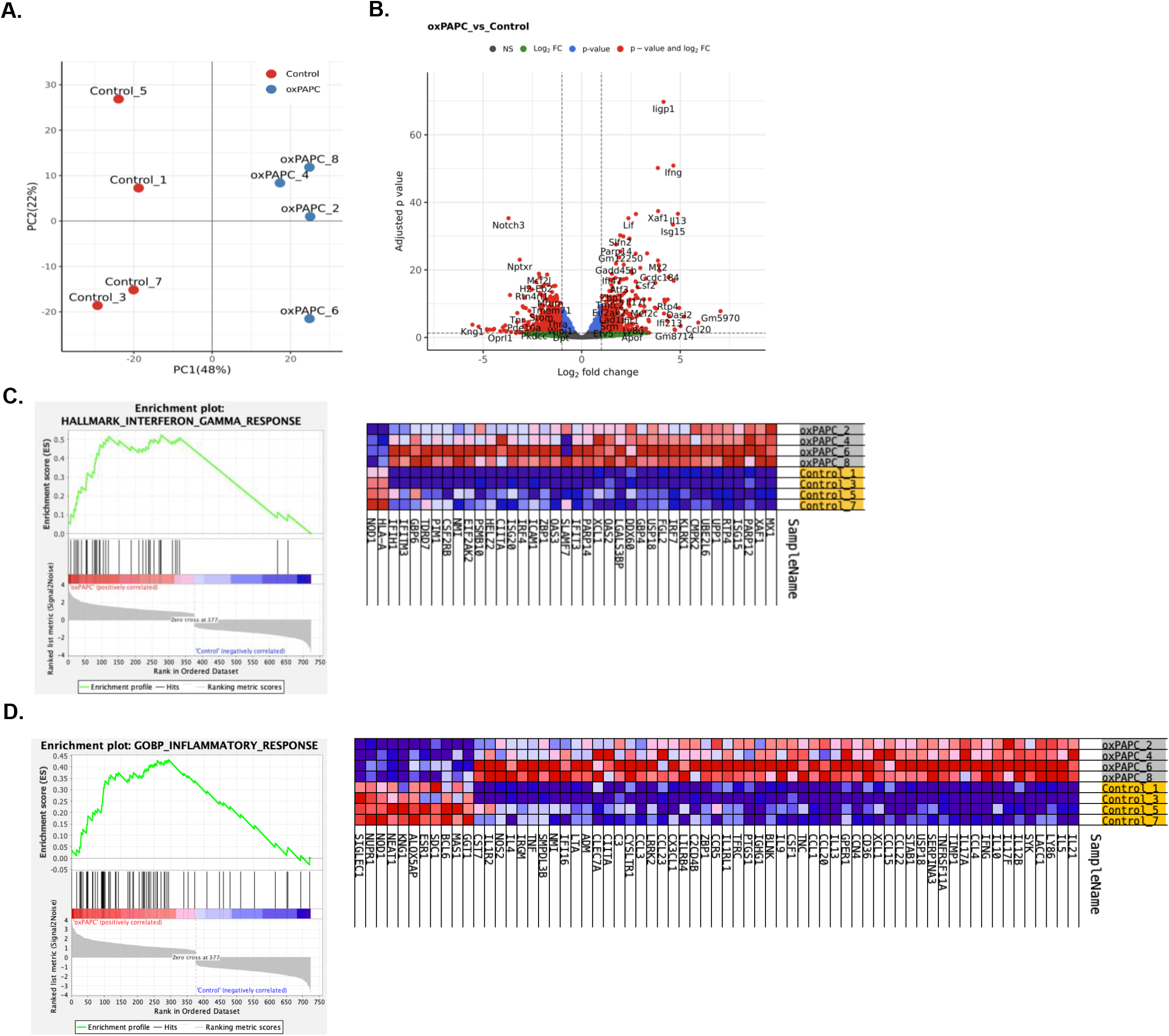
OxPAPC treated T_regs_ increase expression of inflammatory and IFN-γ response genes. CD4^+^ T cells were enriched from the spleens of four- to six-week-old B6 mice and skewed to a T_reg_ phenotype for three days with or without 5μg/ml oxPAPC before being harvested for bulk RNA sequencing. N=4 mice per group. Shown is one of two independent experiments. **(A)** Principal component analysis comparing control and oxPAPC-treated T_regs_. **(B)** Volcano plot showing differentially expressed genes in oxPAPC-treated T_regs_ (significantly enriched = red and right, significantly reduced = red and left). **(C)** GSEA results and heatmap for significantly altered genes in IFN-γ response pathway (red=high, blue= low). **(D)** GSEA results and heatmap for significantly altered genes in inflammatory response pathway (red=high, blue= low).

**Figure 3:**
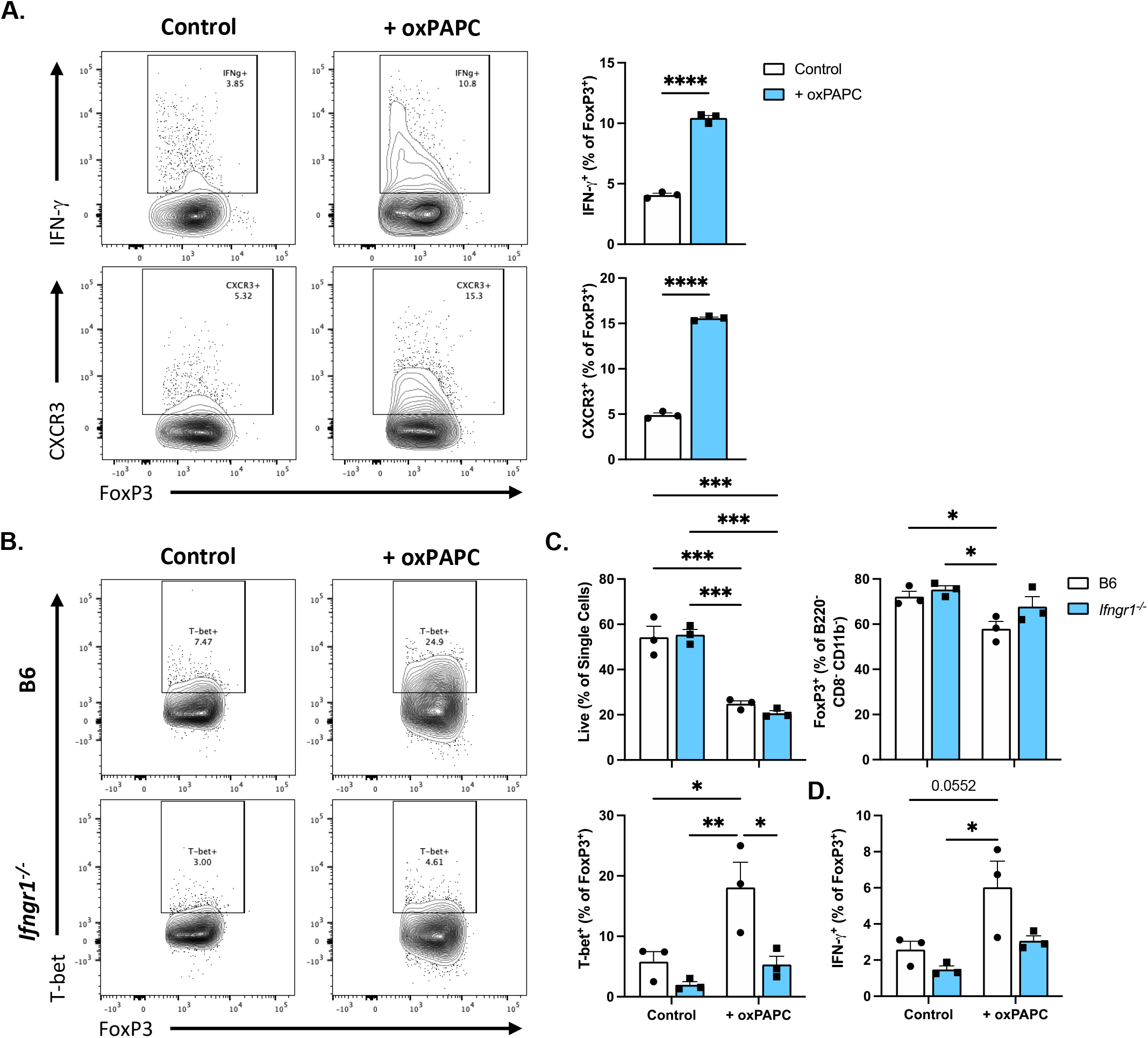
OxPAPC treatment increases T_reg_ IFN-γ production through an IFNγR1-dependent mechanism. **(A)** CD4^+^ T cells were enriched from the spleens of four- to six-week-old B6 mice and skewed to a T_reg_ phenotype for five days with or without 5μg/ml oxPAPC. Th1 associated cytokines and markers were assessed by flow cytometry after a four-hour stimulation with PMA, ionomycin, and Golgi inhibitor (top) or directly after harvest (bottom). **** denote significance *p*<0.0001 by Student’s *t* test. CD4^+^ T cells were enriched from the spleens of four- to six-week-old B6 or *Ifngr1*^*-/-*^ mice and skewed to a T_reg_ phenotype for three **(B**,**C)** or five **(D)** days with or without 5μg/ml oxPAPC. T_reg_ phenotype was assessed by flow cytometry directly at harvest **(B**,**C)** or after a four-hour stimulation with PMA, ionomycin, and Golgi inhibitor **(D)**. Shown is one representative of at least three experiments. Data points are individual mice and error bars show SEM. *, **, and *** denote significance *p*<0.05, *p*<0.01, and *p*<0.001, respectively, by two-way ANOVA and Tukey’s multiple comparison test.

### OxPAPC-treated T_regs_ have an IFN-γ enriched genetic signature

Bulk RNAseq was performed on T_regs_ after three days in culture in the presence or absence of oxPAPC. Distinct transcriptional profiles were observed between the two groups (Figure 2A) and differential expression analysis identified *Ifng* as the second highest increased gene (Figure 2B). Furthermore, gene set enrichment analysis (GSEA) revealed increased expression of genes in the IFN-γ response pathway (Figure 2C) as well as the inflammatory response pathway (Figure 2D) in oxPAPC-treated T_regs._ Together these results indicate oxPAPC treatment induces widespread inflammatory changes in the T_reg_ transcriptome identifying response to IFN-γ as a potential mechanism.

### OxPAPC induces a Th1-like phenotype in T_regs_

Given the IFN-γ signature identified in oxPAPC-treated T_regs_, we sought to investigate the expression of the cytokine and other Th1-associated markers in our oxPAPC-treated T_reg_ cultures. Consistent with RNAseq, the proportion of IFN-γ^+^ FoxP3^+^ cells and CXCR3^+^ FoxP3^+^ cells in oxPAPC-treated cultures were significantly increased (Figure 3A). The *in vitro* T_reg_ phenotype induced by oxPAPC was consistent with *in vivo* T_regs_ in atherosclerotic mice. We compared T_regs_ from male *Ldlr*^*-/-*^ mice fed a high fat Western diet (WD) or normal chow (NC) (Supplemental Figure 4A). In accord with previous studies^23,33^ and our *in vitro* results, WD-fed animals had an increased proportion of IFN-γ expressing T_regs_ in the aorta (Supplemental Figures 4B and 5A), while the proportion of IFN-γ^+^ T_regs_ in the spleen of these mice remained unchanged (Supplemental Figures 4C and 5B). Similar results were observed for NC- and WD-fed female *Ldlr*^*-/-*^ mice (data not shown). Collectively, these data support a role for oxPLs in dysregulation of T_regs_ during differentiation in atherosclerosis.

**Figure 4:**
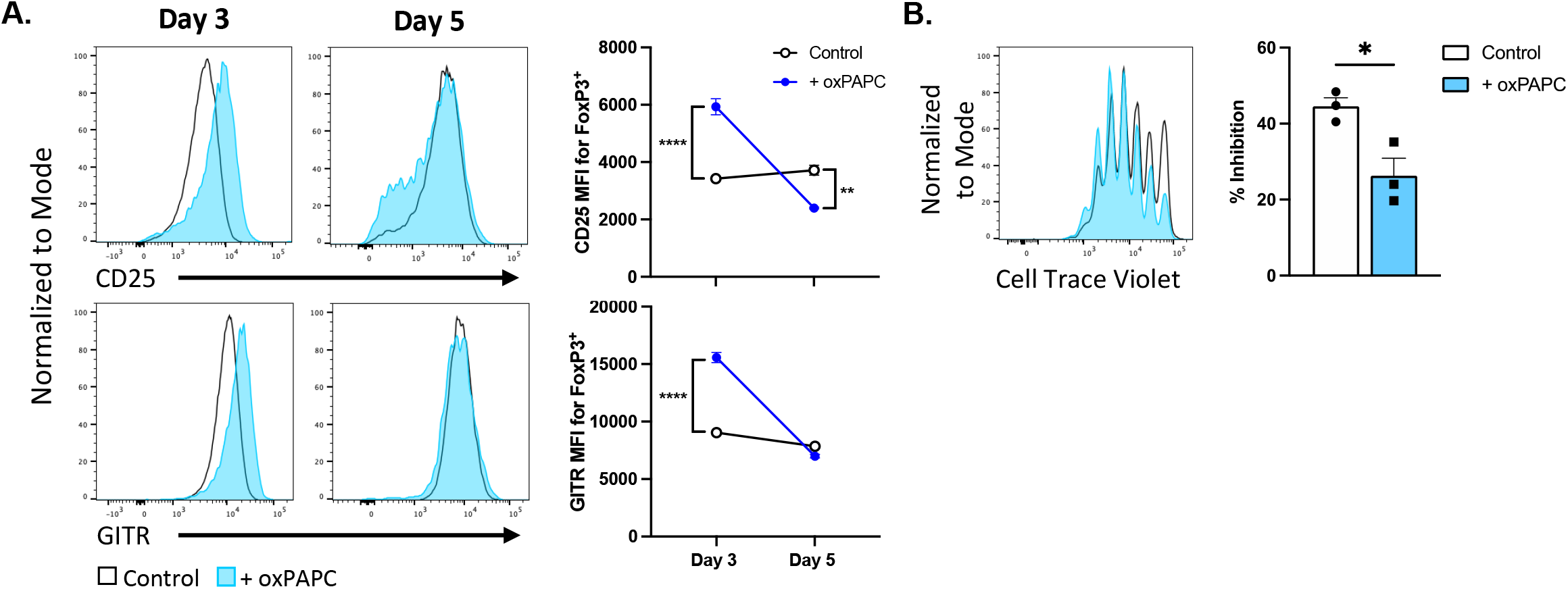
OxPAPC treated T_regs_ are less suppressive. **(A)** CD4^+^ T cells were enriched from the spleens of four- to six-week-old B6 mice and skewed to a T_reg_ phenotype for the indicated time with or without 5μg/ml oxPAPC. T_reg_ markers were assessed by flow cytometry. Error bars show SEM. ** and **** denote significance *p*<0.01 and *p*<0.0001, respectively, by two-way ANOVA and Tukey’s multiple comparison test. **(B)** CD4^+^ T cells were enriched from the spleens of four- to six-week-old B6-Thy1.1-FoxP3^GFP^ mice and skewed to a T_reg_ phenotype for five days with or without 5μg/ml oxPAPC. T_regs_ were then co-cultured with cell trace violet-labeled CD8^+^ T cells for three days. Percent inhibition of CD8^+^ T cell proliferation was assessed by flow cytometry. Shown are the 3:1 T_reg_:T_res_ ratio results from one representative of at least three experiments. Error bars show SEM. * denote significance *p*<0.05 by Student’s *t* test.

### IFN-γ signaling is required for the Th1-like phenotype of oxPAPC-treated T_regs_

Due to the elevated production of IFN-γ by oxPAPC-treated T_regs_ and the known autocrine effects of the cytokine, we tested the contribution of IFN-γ signaling to oxPAPC-mediated T_reg_ phenotypes. To do this, naïve CD4^+^ T cells were purified from spleens of *Ifngr1*^*-/-*^ mice and skewed to T_regs_ in the presence or absence of oxPAPC. Interestingly, the Th1-like phenotype characterized by increased T-bet^+^ IFN-γ producing FoxP3^+^ cells, did not develop in IFNγR1-deficient oxPAPC-treated T_regs_ (Figure 3B-D). However, IFNγR1 deficiency did not affect the oxPAPC-induced reduction in viability (Figure 3C), suggesting cell death is mediated by a separate mechanism.

### OxPAPC reduces T_reg_ suppressive capacity

Because T_regs_ have been shown to be impaired in the oxidized lipid-rich microenvironment during atherosclerosis^20–23^, we hypothesized oxPAPC treatment would make T_regs_ less suppressive. Interestingly, oxPAPC-treated T_regs_ expressed higher levels of functional T_reg_ markers at day 3 in culture, but by day 5 these were significantly reduced, in some cases to below control levels (Figure 4A). In agreement with these observations, when T_regs_ were cultured for five days and then used for an *in vitro* suppression assay, oxPAPC-treated T_regs_ were less able to suppress CD8^+^ T cell proliferation than control T_regs_ (Figure 4B), indicating oxPAPC treatment impairs T_reg_ function as well as altering phenotype. These data suggest that presence of oxPL during induction of T_regs_ decreases their stability over time and thus would affect their suppressive function during atherosclerosis.

To assess the function of oxPAPC-induced T_regs_ in atherosclerosis, T_regs_ were skewed from Thy1.1^+^ mice in the presence or absence of oxPAPC and adoptively transferred into *Ldlr*^*-/-*^ mice with established lesions (Figure 5A). Saline injected *Ldlr*^*-/-*^ mice served as the control. Compared to the saline injected group, mice given control T_regs_ had significantly less atherosclerosis in the proximal aorta. In contrast, oxPAPC-treated T_regs_ were unable to inhibit atherosclerosis progression (Figure 5B-C), demonstrating oxPAPC-induced Th1-like T_regs_ are not protective *in vivo*. Adoptive transfer of T_regs_ did not significantly alter serum cholesterol or triglycerides (Supplemental Figure 6A), indicating the changes observed in the plaque were immune mediated rather than a consequence of improved dyslipidemia. Additionally, oxLDL-specific antibody titers were similar among all groups (Supplemental Figure 6B).

**Figure 5:**
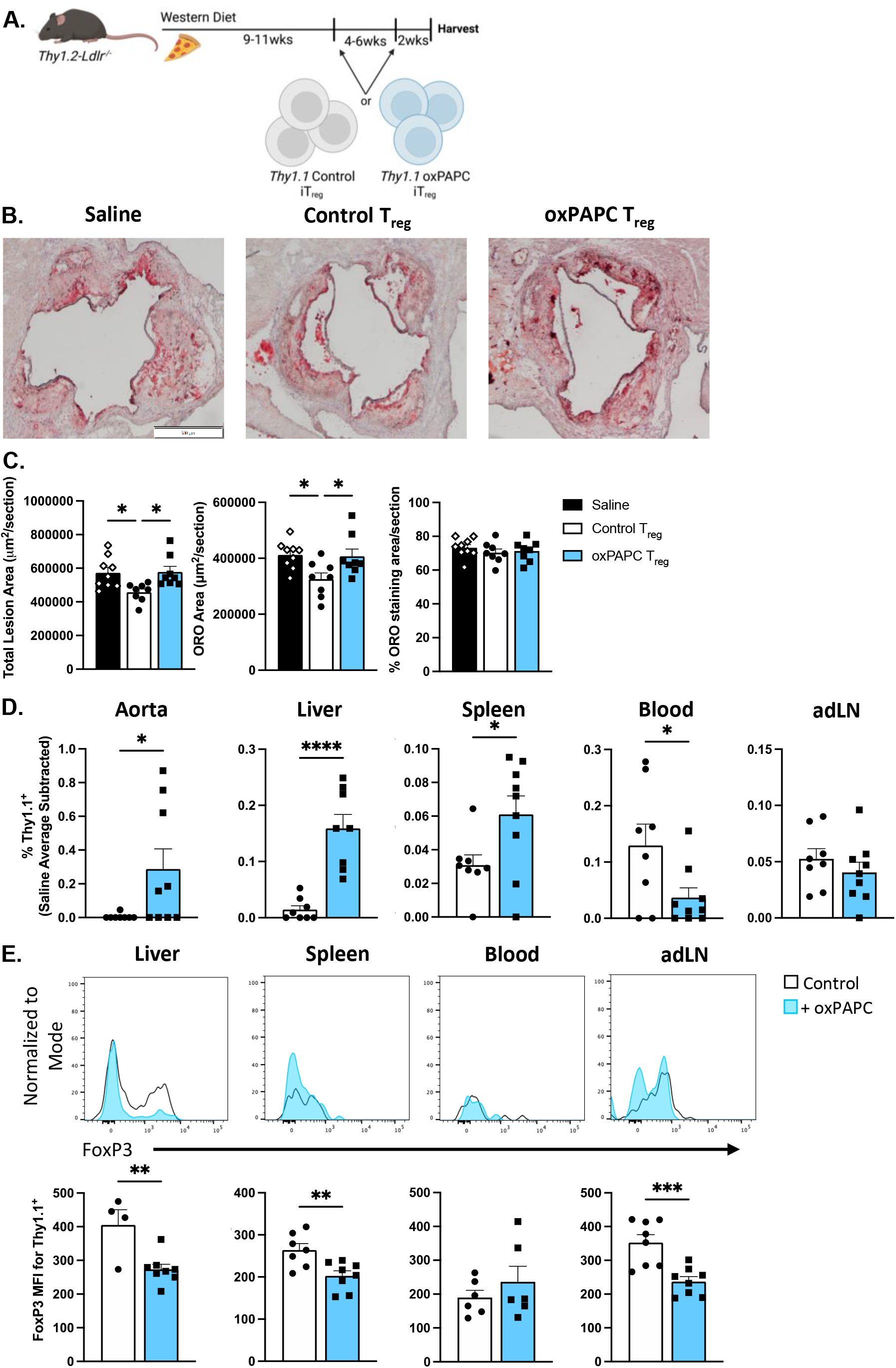
OxPAPC treated T_regs_ are unable to inhibit atherosclerosis progression. **(A)** Study design (image generated with Biorender). Adoptively transferred T_regs_ were skewed from CD4^+^ T cells enriched from the spleens of four- to six-week-old B6-Thy1.1-FoxP3^GFP^ mice for five days with or without 5μg/ml oxPAPC. *Ldlr*^*-/-*^ mice were injected as shown with saline, control T_regs_ or oxPAPC-treated T_regs_. **(B)** Representative Oil-Red-O (ORO) stained atherosclerotic lesions from the aortic root (scale bar represents 500μm). **(C)** Quantification of ORO stained lesions by total lesion area, ORO staining area, and percentage of ORO staining in recipient *Ldlr*^*-/-*^ mice. N=8-9 mice per group, 4 independent experiments. *, **, ***, and **** denote significance *p*<0.05, *p*<0.01, *p*<0.001, and *p*<0.0001 by one-way ANOVA and Tukey’s multiple comparison test. **(D)** Indicated tissues were processed and analyzed by flow cytometry. The average Thy1.1^+^ signal in saline injected controls was treated as background and subtracted from samples. **(E)** Adoptively transferred T_regs_ in the indicated tissues were analyzed by flow cytometry. Histogram shows representative samples. *, **, ***, and **** denote significance *p*<0.05, *p*<0.01, *p*<0.001, and *p*<0.0001 by Student’s *t* test. N=8-9 mice per group, 4 independent experiments. Data points represent individual mice and error bars show SEM.

Thy1.1^+^ cells were detected in all tissues tested from recipient mice (Figure 5D and Supplemental Figure 7A-C). Mice injected with oxPAPC-treated T_regs_ had a larger proportion of transferred cells in the aorta, liver, and spleen, but a significant reduction circulating in the blood compared to control T_reg_ recipients (Figure 5D). Both control and oxPAPC-treated T_regs_ had reduced FoxP3 expression relative to when they were first injected, but adoptively transferred oxPAPC-induced T_regs_ in the liver, spleen and adLNs expressed significantly less FoxP3 than control T_regs_ by the end of the study (Figure 5E). Interestingly, this relative reduction in FoxP3 was not observed in transferred cells found in the blood (Figure 5E). Overall, these data indicate that oxPAPC not only alters T_reg_ phenotype, but also compromises their atheroprotective function, possibly by reducing T_reg_ stability and altering migration *in vivo*.

## Discussion

The current study demonstrates that oxPAPC elicits a Th1-like phenotype in T_regs_ and compromises their function *in vitro* and *in vivo*. Previous work has shown T_regs_ are reduced in number and are less suppressive after treatment with oxLDL^15,25^, in agreement with our own findings with oxPAPC. However, these prior studies utilized fully differentiated T_regs_ for their experiments meaning they were not able to address what role oxLDL might play during T_reg_ polarization. Gaddis *et al*. skewed naïve *Apoe*^*-/-*^ CD4^+^ T cells *in vitro* with TGF-β and reported the inclusion of oxLDL in these cultures inhibited T_reg_ differentiation^22^. However, oxLDL is a rather large, heterogeneous molecule, so this work does not specifically address the role of oxPL in T_reg_ dysregulation. Therefore, our work fills a gap in knowledge regarding how oxPLs, as represented by oxPAPC, impact the differentiation and function of T_regs_.

It is known that adoptive transfer of T_regs_ into atherogenic mouse models inhibits plaque progression^3,14,15,36^. This is consistent with our current findings which showed transfer of control T_regs_ into Western diet-fed *Ldlr*^*-/-*^ mice significantly reduced lesion size. However, the adoptive transfer of oxPAPC-treated T_regs_ did not result in reduced atherosclerosis suggesting exposure to oxPLs reduce the suppressive capacity of T_regs_ *in vivo* (Figure 5). Transferred T_regs_ also demonstrated differential migration in hyperlipidemic mice. In oxPAPC-treated T_reg_ recipients, cells migrated to the tissues (aorta, liver, spleen), whereas the control T_regs_ tended to remain in the blood (Figure 5). These findings are consistent with work from Amersfoort *et al*. showing T_regs_ from dyslipidemic mice homed to atherosclerotic lesions and sites of inflammation^37^. Altogether these results indicate oxPLs directly promote functional and migratory changes in T_regs_ that ultimately undermine their regulatory function and allow for disease progression.

T_reg_ instability is one major hypothesis for T_reg_ loss in atherosclerosis with exT_regs_ developing Th1- or T follicular helper-like pathogenic phenotypes^22,23^. Here, we demonstrate that oxPAPC treatment leads to an overall loss of FoxP3 expression over time in culture. Consistent with potential T_reg_ instability, Th1-like markers (T-bet, IFN-γ, CXCR3) were increased in FoxP3^lo^ T_regs_, a population that proportionally increased with oxPAPC treatment, while receptors related to T_reg_ function (CD25, GITR, ICOS) associated with FoxP3^hi^ cells (Supplemental Figure 3). Additionally, oxPAPC-treated T_regs_ had reduced expression of FoxP3 compared to control T_regs_ following adoptive transfer into hyperlipidemic *Ldlr*^*-/-*^ mice (Figure 5). Collectively, this suggests oxPLs, like oxPAPC, may be partly responsible for T_reg_ instability in the plaque. This conclusion is consistent with the findings of Wolf *et al*.^38^ and Kimura *et al*.^39^ which show T_regs_ specific for the apolipoprotein B core of oxLDL have increased expression of T effector-associated markers in mice and humans with atherosclerosis.

The potential for oxPL-induced T_reg_ instability also has implications for atherosclerosis regression. In Sharma *et al*. T_regs_ in progressing atherosclerotic plaques are mostly thymically derived T_regs_ characterized by Nrp1 expression. However, regressing plaques, in mice where lipid levels were normalized, have a larger T_reg_ population in part due to newly arrived Nrp1^-^ inducible T_regs_ from the periphery^17^. These findings suggest the normalization of oxPLs in circulation allows for altered recruitment or survival of peripherally induced T_reg_ populations. Additionally, Overacre-Delgoffe *et al*. have shown that in the context of the tumor microenvironment, T_regs_ lacking Nrp1 are sensitive to IFN-γ induced dysfunction^40^. These works, in combination with our own findings showing an IFN-γ dependent mechanism for oxPAPC-induced Th1-like T_regs_ (Figure 3), suggest a model in which oxPLs induce T_reg_ instability in peripherally derived Nrp1^-^ T_regs_. This may lead one to hypothesize that peripherally induced Nrp1^-^ T_regs_ differentiating *in situ* or migrating to the plaque during disease progression cannot persist as functional T_regs_ due to increased presence of oxPLs.

In addition to T_reg_ instability, T_reg_ numbers are also reduced during atherosclerosis as a result of increased apoptosis^20,21^. Magnato-García *et al*. proposed that the increase in T_reg_ death is due to the altered microenvironment of the plaque^20^ and others have shown that oxLDL is capable of inducing apoptosis in T_regs_^21,25^. Our results demonstrating oxPAPC-treated T_regs_ undergo significantly increased levels of apoptosis compared to control T_regs_ (Figure 1 and Supplemental Figure 2), are consistent with these findings and indicate oxPLs are at least one specific component of the lesion microenvironment that may induce T_reg_ apoptosis. Interestingly, oxPAPC-induced T_reg_ apoptosis was not mediated by IFNγR1 signaling unlike the Th1-like phenotypes (Figure 3). T_reg_ death was also not due to an overall toxicity of oxPAPC as neither Th1 nor Th17 cells polarized with oxPAPC had reduced viability (Figure 1). OxPAPC can bind to several T cell-expressed receptors including TLR2 and 4, and CD36. We hypothesize oxPAPC signaling through TLR-4 is responsible for the observed cell death as it has been shown to mediate apoptotic signals in other cell types^41–44^, but this will require formal testing in future studies.

Th1-like T_regs_ such as those we describe here, are not specific to atherosclerosis and have been identified in the literature in various inflammatory conditions. Interestingly, in atherosclerosis, cancer, and vitiligo these cells are described as proinflammatory^23,33,40,45^, while in graft-versus-host disease, diabetes, and infection they are anti-inflammatory^46–49^. Therefore, our findings may suggest the ultimate impact of these Th1-like T_regs_ is disease dependent and perhaps in niches characterized by high levels of oxPL, like atherosclerotic plaques and tumors, Th1-like T_regs_ are dysfunctional. This is consistent with Li *et al*. proposed importance for unidentified “atherosclerosis antigens” to the development of a proinflammatory Th1-like T_reg_ phenotype^33^.

In conclusion, our study shows oxPAPC elicits a Th1-like phenotype specifically in T_regs_, compromising their suppressive function. Differentiation of Th1-like T_regs_ is dependent on IFN-γ signaling, but a yet to be identified pathway exists to mediate oxPAPC-induced T_reg_ apoptosis. Overall, our findings demonstrate that oxPLs can directly alter T_reg_ phenotype and may be relevant to their decreased presence and function in atherosclerosis.

## Acknowledgements

We would like to thank Jeffrey C. Rathmell for use of his MacsQuant Analyzer and Vanderbilt Technologies for Advanced Genomics Analysis and Research Design (VANGARD) for assistance with analysis of the RNA sequencing data.

## Sources of Funding

This work is supported by grants from the National Institutes of Health (R01AI153167 to A.S. Major; F31HL154569 to B.D. Appleton; 5T32AR059039-07 to B.D. Appleton), American Heart Association (20IPA35380020 to A.S. Major), and the Veterans Association (VA Merit I01BX002968 to A.S. Major).

## Disclosures

None.

## Notes

### Competing Interest Statement

The authors have declared no competing interest.

